# Coupling MALDI-TOF mass spectrometry protein and specialized metabolite analyses to rapidly discriminate bacterial function

**DOI:** 10.1101/215350

**Authors:** Chase M. Clark, Maria S. Costa, Laura M. Sanchez, Brian T. Murphy

**Affiliations:** Department of Medicinal Chemistry and Pharmacognosy, College of Pharmacy, University of Illinois at Chicago, 833 South Wood Street (MC 781), Room 539, Chicago, IL, United States; Faculty of Pharmaceutical Sciences, University of Iceland, Hagi, Hofsvallagata 53, IS-107 Reykjavík, Iceland

**Author notes:** C.C and S.C. contributed equally to this work. L.S. and B.M jointly supervised this work. Correspondence and requests for materials should be sent to B.M. or L.S.

**Keywords:** natural products, MALDI-TOF MS, bioinformatics, specialized metabolism

## Abstract

For decades, researchers have lacked the ability to *rapidly* correlate microbial identity with bacterial metabolism. Since specialized metabolites are critical to bacterial function and survival in the environment, we designed a data acquisition and bioinformatics technique (IDBac) that utilizes *in situ* matrix-assisted laser desorption/ionization time-of-flight mass spectrometry (MALDI-TOF MS) to analyze protein and specialized metabolite spectra of single bacterial colonies from agar plates. We demonstrated the power of our approach by discriminating between two *Bacillus subtilis* colonies in under 30 minutes, which differ by a single genomic mutation, solely on the basis of their differential ability to produce cyclic peptide antibiotics surfactin and plipastatin. Next, we employed our IDBac technique to detect subtle intra-species differences in the production of metal scavenging acyl-desferrioxamines in a group of eight freshwater *Micromonospora* isolates that share >99% sequence similarity in the 16S rRNA gene. Finally, we employed our method to simultaneously extract protein and specialized metabolite MS profiles from unidentified species of Lake Michigan sponge-associated bacteria cultivated on an agar plate. In just 3 hours, we created hierarchical protein MS groupings of 11 environmental isolates (10 MS replicates each, for a total of 110 samples) that accurately mirrored phylogenetic groupings. We further distinguished isolates within these groupings, which share nearly identical 16S rRNA gene sequence identity, based on inter- and intra-species differences in specialized metabolite production. To our knowledge, IDBac is the first attempt to couple *in situ* MS analyses of protein content *and* specialized metabolite production to allow the distinction of closely related bacterial colonies.

**Significance:** Mass spectrometry is a powerful technique that has been used to identify bacteria via protein content, and to assess bacterial function in an environment via analysis of specialized metabolites. However, until now these analyses have operated independently, and this has resulted in the inability to rapidly connect bacterial phylogenetic identity with patterns of specialized metabolism. To bridge this gap, we designed a MALDI-TOF mass spectrometry data acquisition and bioinformatics pipeline (IDBac) to discriminate both intact protein and specialized metabolite spectra directly from bacterial cells grown on agar. To our knowledge, this is the first technique that organizes bacteria into highly similar phylogenetic groups and allows for comparison of metabolic differences of hundreds of isolates in just a few hours.

---

For nearly two centuries researchers have studied bacteria to diagnose and treat diseases, elucidate intricate inter- and intra-species evolutionary processes, manage and develop agricultural biocontrol practices, and broadly speaking, learn about the complex roles of microorganisms in the environment. Thus, developing techniques to rapidly characterize and discriminate between bacteria has been paramount to these efforts. In the past four decades, sequencing of the 16S ribosomal RNA (rRNA) gene has been instrumental to the classification of bacteria due to its widespread presence in the Kingdom, degree of conservation, and size (1, 2). This and other genetic-based approaches such as pulsed field gel electrophoresis, multilocus sequence typing, and DNA-DNA hybridization have become commonplace (3). However, several limitations to these techniques including cost, turnaround time needed for sequencing and/or analysis, narrow windows of “universal” primers, and in some cases low species-level phylogenetic resolution, have limited their application. Most importantly, in the majority of cases these methods are unable to elucidate how microorganisms interact with one another and function *in situ*. To address this shortcoming, we developed a pipeline that allows rapid discrimination of bacteria based on mass spectral signatures of functional specialized metabolite production in complement to conserved ribosomal housekeeping proteins. This technique is particularly useful for using specialized metabolite production to distinguish strains that share >99% 16S rRNA gene sequence identity.

Specialized metabolite production represents functional traits in bacteria; this principle is exemplified in studies involving the marine obligate bacterial genus *Salinispora* (4–6). To date, the suite of molecules described from *Salinispora* has served to complement existing phylogenetic methods to more precisely differentiate closely related species (7). For example, extensive population-level biosynthetic gene cluster (BGC) diversity was found within three closely related species that shared 99% 16S rRNA gene sequence identity (*S. arenicola, S. pacifica, S. tropica*) (5). Of 75 sequenced genomes within this group of three species, a surprising 229 distinct polyketide synthase (PKS) and nonribosomal peptide synthetase (NRPS) operational biosynthetic units (OBUs) were predicted, supporting that in some cases the acquisition of biosynthetic machinery to produce specialized metabolites is reliant upon a strain’s surrounding environmental pressures. These findings highlight the limitations of employing phylogenetic approaches based on 16S rRNA genes to infer bacterial function.

Shortly after the development of 16S rRNA gene sequencing, MALDI-TOF MS was implemented as a technique to identify large biomolecules (8, 9). Subsequent innovations in instrumentation led to the ability to obtain better resolved spectra of intact proteins in high throughput, facilitating the rapid and less-costly identification of bacteria based largely on ribosomal MS fingerprints (9–12). Bruker (13) and bioMerieux (14) have successfully applied this technology in the clinical setting, while many others have employed it on relatively small strain groupings (from ten to a few hundred isolates) in a genus and species-specific manner to environmental bacteria, or to the classification of mammalian cells, as recently summarized by these reviews (15–17). While MALDI MS protein profiling is useful to determine the putative genus and species-level groupings of bacterial isolates, (18, 19) it does not provide information on their specialized metabolites, which are critical to survival and adaption to the surrounding environment. Relatively little is known about the relationship between bacterial taxonomy and specialized metabolite production in the majority of bacteria isolated from the environment, yet these characteristics are central to researchers who study bacteria in both academic and industrial settings. Thus, a comprehensive analytical pipeline that allows simultaneous analyses of these factors has been a major obstacle to correlating microbial identity with specialized metabolite production (*e.g*. bacterial functional traits).

In this study we present a significant innovation to previously described MS methods that analyze bacteria by utilizing the full capabilities offered by MALDI-TOF mass spectrometers. In addition to linear mode protein analysis, we employ reflectron mode to analyze specialized metabolites, as the combination of both information-rich spectral regions has yet to be applied to existing MALDI-TOF MS analysis pipelines. Silva et al. recently provided a comprehensive history detailing the relatively limited use of MALDI MS to analyze specialized metabolites(20). Utilizing reflectron mode provides increased resolving power and mass accuracy that allows us to inventory bacterial specialized metabolite production in seconds.

We employ consecutive protein and specialized metabolite MALDI-TOF MS analyses on cellular material scraped from single colonies of bacteria grown on agar plates in order to rapidly discriminate intra-species differences in functional chemistry, often on colonies that share indistinguishable colony morphology. To validate our pipeline, we demonstrate the ability to differentiate isolates within closely related *Bacillus*, *Paenibacillus*, and *Micromonospora* species groupings and characterize them based on *in situ* antibiotic, siderophore, and motility factor production. Total acquisition, analysis, and visualization of MALDI-TOF MS data from *both* intact proteins and specialized metabolites of up to 384 bacterial colonies grown on agar can be performed in less than 4 hours and requires minimal MS expertise, compared to that required to operate qTOF or FTCIR mass spectrometers. This provides an alternative to laborious liquid cultivation, metabolite extraction, and chromatographic experiments that are the current standard of practice for studies that focus on specialized metabolite analysis. To our knowledge IDBac is the first attempt to couple *in situ* MS analyses of protein content and specialized metabolite production to afford detailed chemical profiles that allow the distinction of closely related bacterial colonies.

## Results

In order to visualize relational patterns between bacterial isolates, we created a bioinformatics pipeline designed to facilitate the multi-stage analysis of protein and specialized metabolite MS data. Generally speaking, we analyzed MS fingerprints of intact proteins (3,000-15,000 Da) and specialized metabolites (200-2,000 Da) in consecutive MALDI-TOF MS linear and reflectron mode acquisitions, respectively. Principal component analysis and hierarchical clustering of protein spectra placed bacterial isolates into putative genus- and species-level groups. Isolates within each grouping were then further discriminated based on differences in specialized metabolite production through analysis of Metabolite Association Networks (MANs) -(see Text S1 for a description). The IDBac software, written in R, may be downloaded and installed via a simple Windows installer (see “Publication Code and Data Availability”). The software was designed for simplicity and ease-of-use, with documentation provided at each step in the workflow which, at the time of publication, provides hierarchical clustering, principle components analysis, and MAN analysis. However, whereas similar software has required users to convert raw data on their own and format these to software-dependent configurations, IDBac processes spectra directly from raw data. This provides a transparent and repeatable data-handling process that is less prone to user-error or data-tampering, and as a byproduct, bundles spectra by sample into the widely accepted and readily-shareable mzXML format. The IDBac software is available under a GNU General Public License, and links to the full code along with data acquisition and analysis tutorials may be found within the “Publication Code and Data Availability” section.

Although our platform is intended to visualize inter- and intra-species differences in specialized metabolite production between colonies rather than to identify the precise chemical structure of excreted specialized metabolites, it was important to validate that specific nodes (*m/z* features) in our MANs represented bacterial chemistry as opposed to matrix peaks, media components, or instrument noise. After binning peaks from our MALDI-TOF MS spectra, we accounted for the matrix and media ionizable compounds by subtracting a reference spectrum for a matrix and media control during each experimental run. To account for instrument noise we precluded signals from the peak picking algorithm that fell below a user-defined signal to noise ratio (*i.e*., 4:1) and retained peaks occurring in a minimum of 70% of replicates. Using the closely related lab-domesticated strains *Bacillus subtilis* 3610 and its frame-shift mutant *B. subtilis* PY79, we demonstrated the ability of IDBac to use specialized metabolite production to distinguish between nearly identical strains based on functional chemistry (Fig. 1). The two strains were chosen based on their relation to one another: *B. subtilis* PY79 (21, 22) is deficient in the ability to produce the antibiotics surfactin and plipastatin due to a frameshift mutation in *sfp*, the 4'-phosphopantetheinyl transferase that mediates non-ribosomal peptide synthetase apoform activation (23).

*B. subtilis* 3610 and *B. subtilis* PY79 were grown on separate A1 nutrient agar plates in replicates of ten, and their specialized metabolite regions were analyzed using MALDI-TOF MS. The differences in specialized metabolite production between strains 3610 and PY79 were readily observed and visualized in our MAN (Fig. 1B), where large nodes represent bacterial colonies and smaller nodes represent *m/z* values (singly charged molecules) present in their corresponding MALDI-TOF spectra. This MAN allowed for facile visualization of both the differences in specialized metabolite spectra between strains and the ions that were shared or unique to each strain. Four plipastatin (**1**-**4**) and three surfactin (**5**-**7**) analogs, including their resolved isotopes and adducts, were detected from strain 3610 and not from PY79, while both strains shared the ability to produce partially characterized polyglutamate polymers, which have been implicated in species-specific functions such as virulence factor production, biofilm formation, and sequestration of toxic metal ions (24, 25). Importantly, the majority of *m/z* values (33/42; 78.6%) in the MAN represented specialized metabolites, and IDBac successfully filtered out ions associated with matrix and media components. MALDI-TOF MS analysis of the specialized metabolite region and data processing using IDBac correctly depicted subspecies antibiotic production differences between genetic variants of *B. subtilis* in under 30 minutes.

**Fig. 1.**
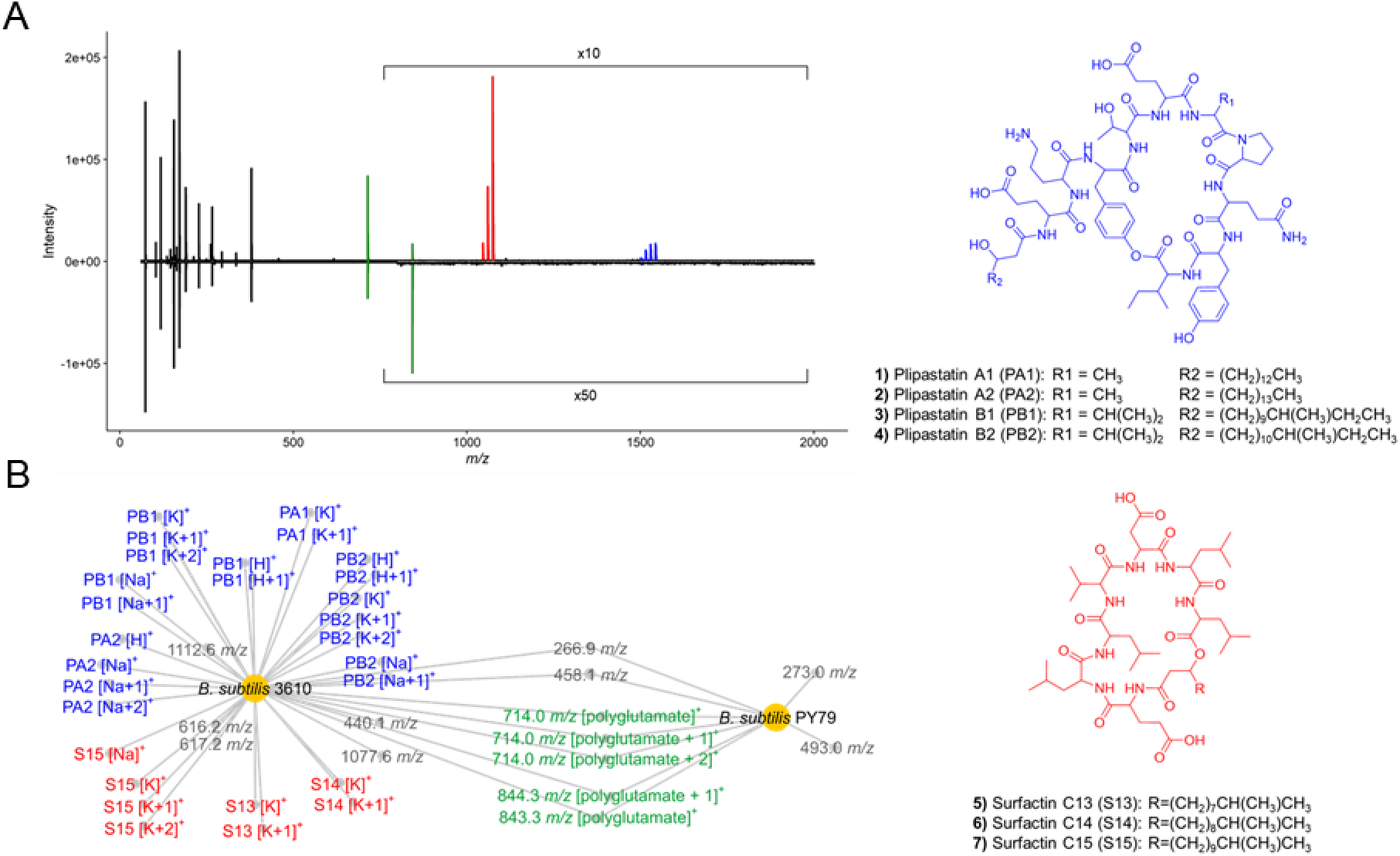
Analysis of MALDI-TOF MS specialized metabolite data from two *B. subtilis* genetic variants (ten replicates each) shows distinct differences in antibiotic production. (A) Inverse spectrum comparison showing representative spectra for *B. subtilis* 3610 (positive spectrum) and *B. subtilis* PY79 (negative spectrum). (B) MAN showing the differential production of surfactin and plipastatin antibiotic analogs between the two strains. Large nodes represent individual bacterial colonies, while smaller nodes represent individual *m/z* values in MALDI-TOF spectra that fall within our peak selection criteria. MALDI-TOF spectrum annotations can be found in Fig. S1. Isotopes are denoted as +1, +2.

Next, we demonstrated that intra-species strain groupings based on MALDI-TOF MS intact protein profiles can be further discriminated based on *in situ* specialized metabolite production. From our in-house strain library, we selected eight *M. chokoriensis* isolates from two sediment samples collected nearly 175 km apart in Lake Michigan, USA. These isolates share >99% 16S rRNA gene sequence identity across >1,460 nucleotides (26). We cultivated biological replicates in a random pattern across a 48-well microwell plate resulting in at least four independent cultures of each strain. With a minimum of eight technical replicates, we used MALDI-TOF MS to consecutively acquire MS spectra of the protein and specialized metabolite regions. This resulted in at least 32 replicate spectra per isolate, with peaks required to be present in 70% or greater of replicates to be included in analyses (a user-defined parameter within IDBac). We observed that hierarchical clustering of MALDI-TOF MS protein data correlated strongly with 16S rRNA similarity (Fig. 2A), results consistent with many previous efforts that employed MALDI-TOF MS as an alternative to traditional sequence-based taxonomic classification methods, as recently summarized (15, 16).

Importantly, while these MALDI-TOF MS protein fingerprints correlated nearly identically with corresponding phylogenetic groupings (Fig. 2A), differences in specialized metabolite production existed and *could not be* predicted through 16S rRNA phylogenetic analyses or geographic strain distribution patterns. However, analysis of IDBac’s MAN rapidly discerned subtle, but significant variations in bacterial chemistry within these closely related *Micromonospora* isolates (Fig. 2B), allowing us to assess the relationship of specialized metabolite production to both strain phylogeny and geographic origin. A major distinguishing pattern exhibited by seven of eight *M. chokoriensis* isolates was a group of features between 650-800 *m/z*. This pattern was characterized by successive 14 Da differences beginning at 673.5 *m/z and* extending to 771.6 *m/z,* along with sodium and potassium adducts of each, which we attributed to analogs differing in the addition/subtraction of methylene groups (see Fig. S2). An outlier strain, B001, did not share these features, indicating differential specialized metabolite production within this group of highly similar strains.

To further validate the differential specialized metabolite production observed in the MAN, we employed HPLC-MS/MS analysis for evaluation with the Global Natural Products Social molecular networking platform (27) and comparative metabolomics XCMS analysis (28). HPLC-MS/MS afforded an orthogonal means of analysis through chromatographic separation, a different mechanism of ionization, and tandem mass data that helped verify relationships between observed metabolites. Using GNPS, XCMS, and theoretical isotope abundance spectra (see Fig. S3-S7), we identified a series of acylated desferrioxamine analogs (**8**-**12**). These belong to a class of siderophores that sequester the essential growth factor ferric iron from the environment. One of these, desferrioxamine B, is used clinically to treat metal poisoning. Interestingly, according to our analyses B001 lacks the capacity to produce these acylated analogs (Fig. 2B), and this was readily highlighted upon analysis of the MAN. We also observed several other desferrioxamine analogs produced by these isolates; links to these data are available in Table S1. Also of note, while the phylogenetic identity of *Micromonospora* isolate B031 is more closely related to B001, it overlaps the remaining six isolates in its ability to produce acylated desferrioxamines, highlighting the importance of our method to provide a means other than phylogeny to distinguish between *in situ* function of similar strains.

The HPLC-MS/MS based observation of strain-specific patterns of siderophore production corroborated our initial MALDI-TOF MS results that showed B001, while able to produce unmodified desferrioxamine B, was deficient in the ability to produce a series of acylated desferrioxamine B analogs (**8**-**12**). This is significant, as B001 was isolated from the same 1 cm^3^ sediment sample as six of the seven strains that produced this compound series, highlighting that phylogenetic groupings and geographic location are not sufficient indicators of specialized metabolite production capacity. Our observations of desferrioxamine biosynthetic pathway promiscuity are consistent with previous studies (29–31). However, each of these studies required extensive genome sequencing and/or liquid fermentation experiments followed by chromatographic analyses, whereas analysis through MALDI-TOF MS/IDBac is able to visualize putative phylogenetic relationships *and* intra-species differences in functional chemistry in a few hours, once each colony appears on a Petri dish.

**Fig. 2.**
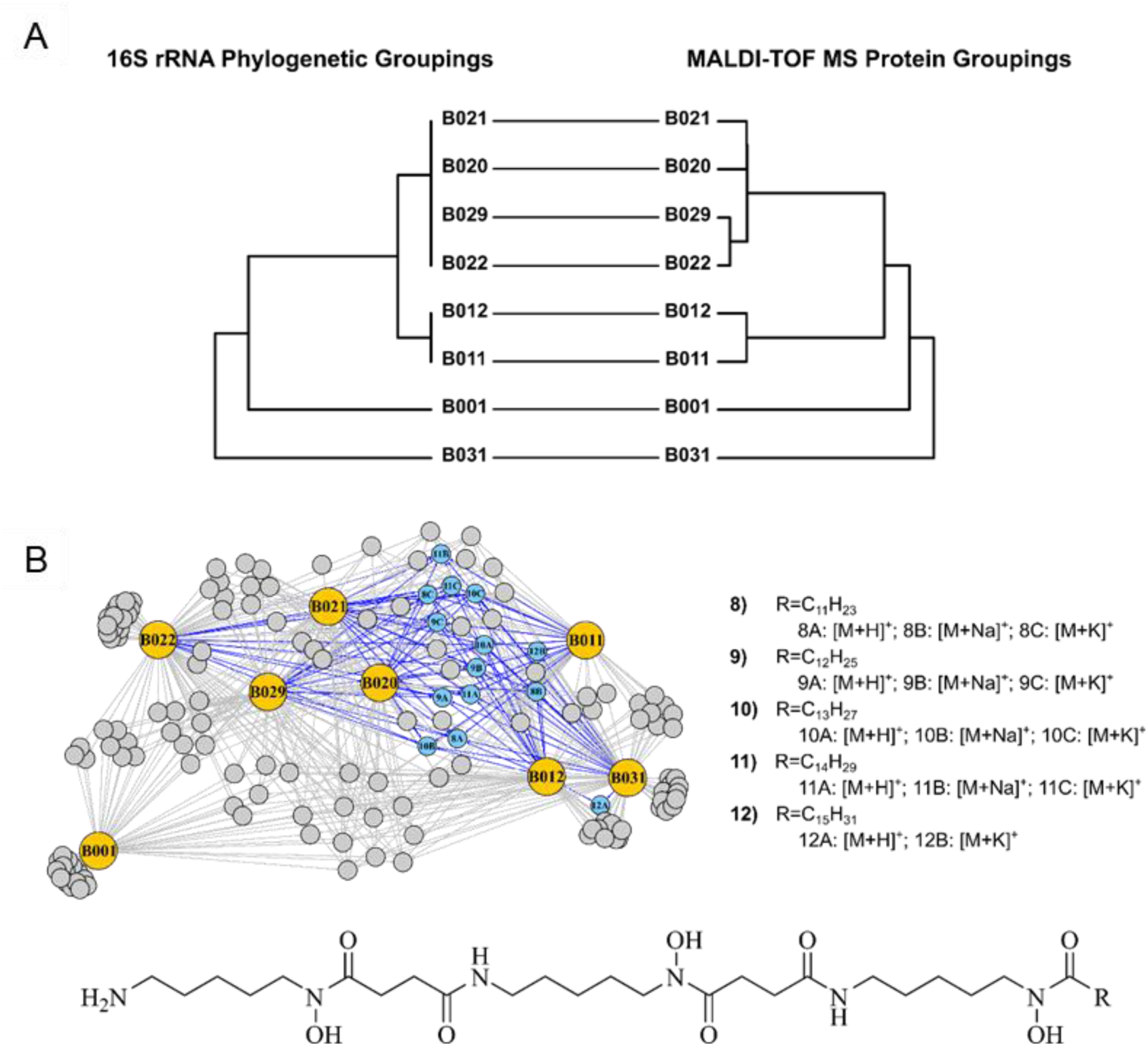
IDBac protein and specialized metabolite analysis of isolates from a single *Micromonospora* species. (A) Tanglegram depicts a high degree of similarity between groupings of 16S rRNA gene sequence identity and MALDI-TOF MS protein data of *M. chokoriensis* isolates. (B) MAN of MALDI-TOF MS data from *M. chokoriensis* colonies highlights distinct intra-species differences in specialized metabolite production; this is due to differential production of a specific series of acylated desferrioxamine siderophores (shown as blue nodes), which B001 did not to produce.

In many instances, the taxonomic identity and specialized metabolite production capacity of a group of bacteria are not well characterized or completely unknown, particularly in studies involving bacteria isolated from the environment. Thus, we tested the ability of the IDBac analysis pipeline to rapidly extract protein and specialized metabolite information from unknown environmental bacteria cultivated from a freshwater sponge collected in Lake Michigan. We cultivated sponge-associated bacteria from 1 cm^3^ of tissue onto high nutrient A1 media (Fig. 3A; see Methods section for further details). Using a sterile toothpick, we selected all colonies that grew on the plate over a 90-day time period. These colonies were then subjected to MALDI-TOF MS analysis in replicates of ten and the data were processed in IDBac. Principal component analysis and unsupervised hierarchical clustering of the bacterial protein range afforded four distinct groupings (Fig. 3B). We confirmed that these MS protein groupings aligned with the genera *Enterococcus*, *Bacillus*, *Paenibacillus*, and *Rhodococcus* via 16S rRNA gene sequencing analysis of each of isolate (Fig. S8-S9).

We next generated a MAN (Fig. 3C) in order to further discriminate these groups based on specialized metabolite production. Interestingly, unsupervised subnetworking modularity analysis (32) separated the network into five groups of strains (colored in Fig. 3C), which highly correlated to phylogenetic groupings. The seven *Rhodococcus* isolates exhibited a high degree of specialized metabolite associations, while the *Bacillus* and *Enterococcus* isolates were classified into unique modules. Importantly, our method quickly highlighted subtle differences in specialized metabolite production between two morphologically identical *Paenibacillus* strains and separated the two strains whose 16S rRNA gene sequences share a pairwise similarity of 99.87% (1496/1498 nucleotides) (26). A careful look at their specialized metabolite profiles shows subtle differences in production of a series of specialized metabolite(s) ranging from *m/z* 900-1250. We wanted to confirm this difference was not due to variations in colony microenvironment, since nearby colonies can affect specialized metabolite production through physical contact and chemical crosstalk (25, 33, 34). To determine this, we grew each *Paenibacillus* isolate by itself over three individual cultivation experiments and MALDI-TOF MS data acquisition events, and after observation of previously observed specialized metabolite patterns in the *m/z* 900-1250 range, we concluded that both isolates contained partially overlapping, but distinct metabolic capacities (see Fig. S10). These subtle differences in metabolic capacity were readily detected and visualized as a result of IDBac specialized metabolite analysis, and could not be achieved through analysis of protein groupings alone. In just three hours our pipeline created statistically robust protein profile groupings of environmental isolates (11 strains, 10 MS replicates each, for a total of 110 MALDI spots). It further distinguished colonies with nearly identical morphology and 16S rRNA gene sequence identity based on their capacity to produce specialized metabolites.

**Fig. 3.**
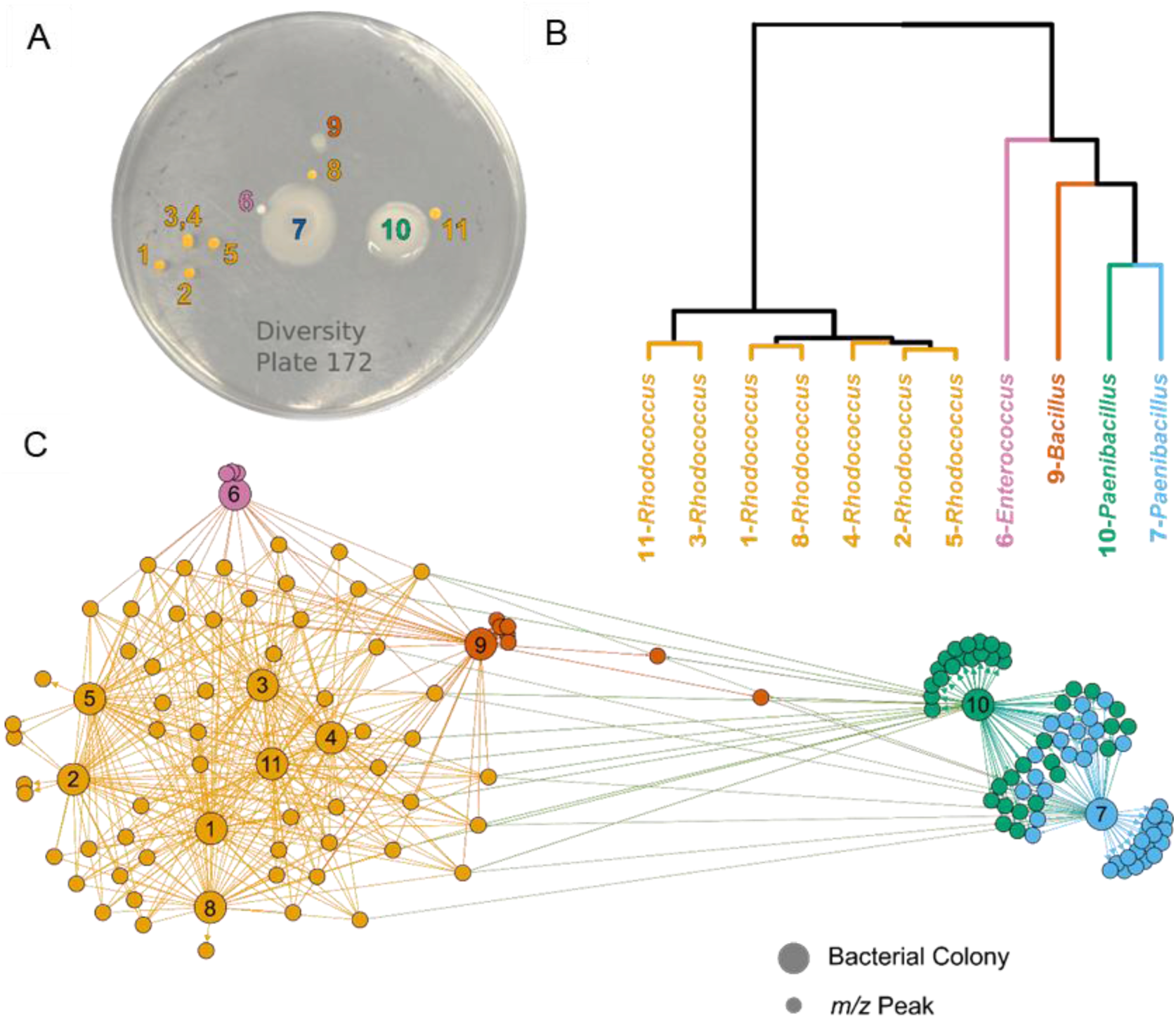
The IDBac pipeline provides rapid protein and specialized metabolite fingerprinting of unknown environmental isolates using MALDI-TOF MS and a freely available bioinformatic interface. (A) Bacterial diversity plate obtained from placing freshwater sponge tissue on high nutrient A1 agar. (B) IDBac allowed for hierarchical clustering of MALDI-TOF MS protein spectra with the option to choose standard distance measures and clustering algorithms. For workflows requiring analysis of hundreds to thousands of strains, protein grouping is essential for data-reduction *before* specialized metabolite analysis is performed. (C) MAN, colored via modularity analysis with default thresholds in Gephi (35), allowed for rapid decision-making based on gross *in situ* specialized metabolite production after matrix and media signals were subtracted automatically from the network in IDBac. *Rhodococcus* isolates shared a common set of core metabolites, while *Enterococcus* and *Bacillus* isolates exhibited less extensive but unique metabolomes. Significant outliers were the two *Paenibacillus* strains, which produced several shared and unique high MW specialized metabolites; this highlights the power of IDBac to visualize intra-species metabolic differences between seemingly identical bacterial colonies, and rapidly empowers the user with an array of taxonomic and metabolic information from which they can generate research questions, form hypotheses, or make informed decisions.

## Discussion

Here we present a method and analysis pipeline to acquire and integrate MALDI-TOF MS data of *both* intact proteins and specialized metabolites from single bacterial colonies. Our technique is unique when compared to other MS-based methods that are commonly employed to analyze specialized metabolites. IDBac rapidly provides a data acquisition and analysis platform to group single bacterial colonies based on putative taxonomic identity, and further discriminates these groups using MS patterns of functional specialized metabolites. No existing MS platform provides the ability to analyze both a diverse array of proteins and non-volatile metabolites at the throughput of hundreds of strains in just a few hours.

One aspect of the IDBac pipeline that provides significant advantages over existing MS platforms is that it affords information on colony taxonomy *and* specialized metabolite production without the need for extraction and chromatographic analyses. The latter techniques rely on growing pure bacterial isolates (often in liquid culture), generating extracts, and using liquid chromatography analyses (generally coupled to MS), to separate and detect specialized metabolites. This is often a laborious, costly, and relatively time-consuming process. Our pipeline is an extraction free process and is ideally suited for researchers that aim to a) compare functional chemistry between closely related isolates (*e.g*, comparing antibiotic production between two *B. subtilis* genetic variants), b) to probe relationships between taxonomic identity and environmental functionality (*e.g*., studying intra-species differences in *M. chokoriensis* siderophore production), or c) to assess the broad relatedness of unknown environmental bacterial isolates (*e.g*., visualizing phylogenetic and metabolic relatedness within a group of unknown bacteria). The latter points highlight a strength of IDBac as an engine to generate research questions and hypotheses. For example, why have similar *Micromonospora* strains from the same 1 cm^3^ of sediment (Table S2) evolved differing capacities to scavenge iron? Is there any correlation between geographic location and desferrioxamine production in an expanded set of Lake Michigan sediment-derived *Micromonospora* isolates? In a Lake Michigan freshwater sponge, is there a synergistic role between the specialized metabolites that are unique to two phylogenetically similar *Paenibacillus* strains? MALDI-TOF MS/IDBac analysis allows for easy visualization of global specialized metabolite patterns, and as a result gives researchers the opportunity to ask these questions based on rapidly acquired data.

IDBac is complementary to other innovative platforms such as GNPS, which aids in the dereplication of previously characterized and identification of potentially new specialized metabolites (27, 36). Successful integration of these orthogonal approaches was demonstrated in our analysis of *M. chokoriensis* isolates. We used IDBac to rapidly identify and discriminate between seemingly identical isolates from a similar environment and detect subtle differences in siderophore production between isolates within the same species grouping. Subsequent growth of these strains, extraction, and LC-MS/MS analyses allowed us to use the GNPS platform to dereplicate the structures of several iron scavenging desferrioxamine analogs. This highlights the individual strengths of orthogonal MS-based approaches to extract complementary information from a biological system.

Through the course of our studies we have documented a few potential limitations of our method. First, although IDBac can create accurate sub-species groupings of bacteria based on protein MS fingerprints, species-level identification is only possible in the presence of a searchable and extensive protein fingerprint database. A greater community effort is required to document MALDI-TOF MS protein fingerprints into a publicly available database to maximize phylogenetic coverage. Such an effort would facilitate the rapid identification of unknown environmental bacteria, and elevate this process to be on par with existing commercial platforms used to identify clinical pathogens (15). An effort to make a readily searchable/freely available bacterial protein MS database is ongoing in our laboratory and the first version of IDBac supports this mission by providing a simple interface for bundling and converting proprietary vendor-format data files to the open-access mzXML format for easy, reproducible sharing.

A second limitation is that bacterial specialized metabolite production is sensitive to external factors such as growth media, temperature, unintended microbial contamination, and the proximity of other microorganisms on a Petri dish (37). For comparison of specialized metabolite profiles to be reliable, it is imperative that strains be cultivated under identical conditions and that several biological/technical replicates are acquired for each isolate (both precautions were taken in this study). Fortunately, the throughput of our pipeline allows for multiple replicates of a sample to be analyzed rapidly. Conversely, MALDI-TOF MS protein fingerprints of bacterial colonies are robust and exhibit minimal fluctuation when the data are acquired on different growth media (Fig. S11).

Finally, our method is limited by the resolution of the MALDI-TOF MS and would be improved through use of instruments with higher resolving power capabilities. High resolution instruments, such as MALDI-FT-ICR, and fragmentation data from a MALDI-TOF/TOF MS could aid in preliminary compound class dereplication via generation of accurate molecular formulas and structural information, respectively. However, this would increase the length of the experiments, data storage requirements, and would require additional technical expertise.

We presented a method that couples bacterial protein and specialized metabolite MS data to rapidly discriminate between isolates based on both their identity and environmental function. Our MS pipeline addresses a need in research communities that study microbial function (*e.g.* chemical ecology, pathogenesis, taxonomy, drug discovery, agriculture and food safety). The pipeline is faster than existing MS methods used to analyze non-volatile metabolite production within microorganisms and requires less technical experience to operate. The experiment requires 10^5^-10^7^ bacterial cells (38) – or approximately the tip of a toothpick – to generate MS profiles, and is designed to be performed in high-throughput. Acquisition of MALDI-TOF MS data from *both* intact proteins and specialized metabolites of single bacterial colonies have been integrated in one single freely available bioinformatics pipeline (IDBac). To our knowledge this is the first report of coupling protein and specialized metabolite MS data in a semi-automated pipeline to make the rapid discrimination of bacteria accessible to the broad research community.

## Methods

### Sponge collection and processing

A freshwater sponge sample was collected June 6th, 2016 from Marinette, Wisconsin (45º5'16.012"N, 87º35'10.468"W), from pilings near Red Arrow Beach at a depth of 8 ft. using SCUBA. The sponge was separated from associated macro-organisms, rinsed with filter sterilized Lake Michigan water five times to remove bacteria from surrounding lake water, and most of the water expelled from the sponge by applying gentle pressure. A 1 cm^3^ section of tissue and 10 mL of sterile 20% glycerol solution were ground for two minutes using an autoclaved mortar and pestle. A 60 ºC dry bath was utilized to pretreat a 500 µL aliquot for 9 minutes. The sample was then diluted 1:10 with 20% sterile glycerol solution. To an agar plate containing A1 nutrient media (5 g of soluble starch, 2 g of yeast extract, 1 g of peptone, 250 mL of filter-sterilized Lake Michigan water and 250 mL of distilled water), 50 µL of sample was added and spread across the surface. The plate was sealed with Parafilm and left at 27 ºC for 90 days.

### MALDI-TOF MS sample preparation

For MALDI-TOF MS analysis, proteins were extracted using an extended direct transfer method that included a formic acid overlay (38). Using a sterile toothpick, bacterial colonies that grew on nutrient agar were applied as a thin film onto a MALDI ground-steel target plate (Bruker Daltonics, Billerica, MA). Over each bacterial smear, 1 µL of 70% LC-MS grade formic acid (Optima, Fisher Chemical) was added and allowed to evaporate, followed by the addition and subsequent evaporation of 1 µL of 10 mg/mL α-cyano-hydroxycinnamic acid (recrystallized from the 98% pure Sigma-Aldrich) solubilized in 50% acetonitrile, 2.5% trifluoroacetic acid and 47.5% water (39). All solvents were HPLC or MS grade.

### MALDI-TOF data acquisition

Measurements were performed using an Autoflex Speed LRF mass spectrometer (Bruker Daltonics) equipped with a smartbeam™-II laser. Detailed instrument settings are available in the Supplementary Information. Specialized metabolite spectra were recorded in positive reflectron mode (5000 shots; RepRate: 2000 Hz; delay: 9297 ns; ion source 1 voltage: 19 kV; ion source 2 voltage: 16.55 kV; lens voltage: 8.3 kV; mass range: 50 Da to 2,700 Da, matrix suppression cutoff: 50 Da). Protein spectra were recorded in positive linear mode (1200 shots; RepRate: 1000; delay: 29793 ns; ion source 1 voltage: 19.5 kV; ion source 2 voltage: 18.2 kV; lens voltage: 7.5 kV; mass range: 1.9 kDa to 2.1 kDa, matrix suppression cutoff: 1.5 kDa). Protein spectra were calibrated externally with the Bruker Daltonics bacterial test standard (BTS). Calibration masses were: RL29, 3637.8 Da; RS32, 5096.8 Da; RS34, 5381.4 Da; RS33meth, 6255.4 Da; RL29, 7274.5 Da; RS19, 10300.1 Da; RNAse A, 13683.2 Da; Myoglobin, 16952.3 Da. Specialized metabolite spectra were calibrated externally using the Bruker Daltonics peptide calibration standard. Calibration masses (monoisotopic) were: CHCA [2M+H]^+^, 379.0930 Da; Angiotensin II, 1046.5418 Da; Angiotensin I, 1296.6848 Da; Substance P, 1347.7354 Da; Bombesin, 1619.8223 Da; ACTH clip 1-17, 2093.0862 Da, ACTH clip 18-39, 2465.1983 Da.

Automated data acquisitions were performed using flexControl software v. 3.4.135.0 (Bruker Daltonics) and flexAnalysis software v. 3.4. Spectra were automatically evaluated during acquisition to determine whether a spectrum was of high enough quality to retain and add to the sum of the sample acquisition. The number of added spectra, quality requirements and other detailed acquisition settings are available in Table S2; for flexControl and flexAnalysis scripts, see the “Publication Code and Data Availability” section.

### 16S rRNA gene sequence analysis and data workup

Bacterial isolates were characterized by analysis of the 16S rRNA gene. Total genomic DNA was extracted using the Ultra Clean Microbial DNA Isolation kit (MOBIO Laboratories) according to the manufacturer’s instructions. The primers 8F, FC27, 1100F, RC1492 and 519R were used for amplifying the 16S rRNA gene. The polymerase chain reaction (PCR) conditions were as follows: initial denaturation at 95°C for 5 minutes, followed by 35 cycles of denaturation at 95°C for 15 seconds, annealing at 60°C for 15 seconds, and extension at 72 °C for 15 seconds, and a final extension step at 72 °C for 2 minutes. PCR products were purified using QIAquick PCR Purification kit from Qiagen and the amplicons sequenced by Sanger sequencing. Geneious V10.0.9 software was used to produce a consensus sequence of the 16S rRNA gene amplified for each bacterial isolate and the identity of the isolates determined using BLASTn. All sequences were submitted to GenBank and accession numbers along with information regarding each of the isolates’ origin is available in Table S2. Phylogenetic trees were created by trimming aligned sequences (SILVA Incremental Aligner)(40) to equal length and using Geneious’ “Tree Builder” with Jukes-Cantor and Neighbor Joining algorithms with 100 bootstrap replicates using a support threshold of 80%.

### Extraction of *Micromonospora* isolates for LC-MS/MS analysis

Extractions were performed from bacterial cultures growing on solid agar media (A1) following the protocol of Bligh, E. G. and Dyer, W.J (41). Agar cultures were divided into 1 cm^3^ pieces and 3 mm glass beads added. Extraction solvent was added in three steps with vigorous vortexing between steps 1) 1:2 (v/v) CHCl_3_:MeOH, 2) CHCl_3_ in 1/3 the added volume of step one, 3) H_2_O in 1/3 the added volume of step one. From the resulting two-layer liquid partition, the organic (bottom) layer was retained for further analysis.

### LC-MS/MS analysis of bacterial extracts

*Micromonospora* extracts were analyzed via LC-MS/MS with a method adapted from that described by Goering et al (42). Experiments were performed on an Agilent 1200 workstation connected to a Thermo Fisher Scientific Q-Exactive mass spectrometer with an electrospray ionization source (ESI). Reversed-phase chromatography was performed by injection of 20 µL of 0.1 mg/mL of extract at a 0.3 mL/min flow rate across a Phenomenex Kinetex C18 RPLC column (150 mm x 2.1 mm i.d., 2 µm particle size). Mobile phase A was water with 0.1% formic acid and mobile phase B was acetonitrile with 0.1% formic acid. Mobile phase B was held at 15% for 1 minute, then adjusted to 95% over 12 minutes, where it was held for 2 minutes, and the system re-equilibrated for 5 minutes. The mass spectrometry parameters were as follows: scan range 200-2000 *m/z*, resolution 35,000, scan rate ∽3.7 per second. Data was gathered in profile and the top 5 most intense ions in each full spectrum were targeted for fragmentation that employed a collision energy setting of 25 eV for Higher-energy Collisional Dissociation (HCD) and isolation window of 2.0 *m/z*. Data were converted to mzXML and uploaded to the GNPS: Global Natural Products Social Molecular Networking platform for dereplication (27) and XCMS (43) for comparative metabolomics. For analysis in R, GNPS library matches were downloaded and converted to mzML, where necessary, with MSConvert. “Amphiphilic ferrioxamine” mzML files were manually edited in Notepad++ to the correct XML schema.

### MALDI-TOF data bioinformatics pipeline

In brief, MALDI-TOF MS raw data were first converted to the open-source mzXML format(44) using ProteoWizard’s command-line MSConvert (45). The mzXML files were read into R through mzr (45) and custom code, and processed into peak lists using MALDIquant (46). Lastly, the peak lists informed interactive data analyses and visualizations that were displayed through a default, offline internet browser with RStudio’s Shiny (47) package. The IDBac software is being distributed under a GNU General Public License, and links to the full code along with data acquisition and analysis tutorials can be found within the “Publication Code and Data Availability” section. Easy to install IDBac software and updates, method manuals and information on code contribution may be found at chasemc.github.io/IDBac and permanent snapshots at https://doi.org/10.5281/zenodo.1115619.

### Publication Code and Data Availability

IDBac software and updates, in addition to method manuals and information on code contribution, may be found at chasemc.github.io/IDBac. The flexControl, flexAnalysis, and R code used to generate all figures and analyses for this publication along with the publication-version of IDBac is available for download at: https://doi.org/10.17632/ysrtr9c5s7.1.

Mass spectra were deposited in MASSIVE for: MALDI

(ftp://massive.ucsd.edu/MSV000081619), and LC-MS/MS

(ftp://massive.ucsd.edu/MSV000081555) data.

## Acknowledgements

The authors wish to acknowledge the following contributors: Russel Cuhel and crew of RV Neeskay at the University of Milwaukee Wisconsin for assistance with sediment collection; Keith Jung for collection of the freshwater sponge; Dr. Amanda Bulman of Bruker for assistance with MALDI-TOF MS protein acquisition parameters; Dr. Terry Moore and Dr. Atul Jain for recrystallizing α-cyano-hydroxycinnamic acid. This publication was made possible by Grant Number K12HD055892 from the National Institute of Child Health and Human Development (NICHD) and the National Institutes of Health Office of Research on Women’s Health (ORWH) (LMS) and UIC startup funds (BTM, LMS).

## Author Contributions

Conceptualization, Methodology & Visualization, B.M., L.S., C.C., S.C.; Software & Validation, C.C.; Investigation & Data Curation, C.C. S.C.; Writing – Original Draft, B.M., C.C.; Writing – Review & Editing, B.M., L.S., C.C., S.C.; Supervision, B.M., L.S.; Project Administration, B.M.

## References

1. Woese CR (1987) Bacterial Evolution. Microbiol Rev 51(2):221–271.

2. Yarza P, et al. (2014) Uniting the Classification of Cultured and Uncultured Bacteria and Archaea Using 16S rRNA Gene Sequences. Nat Rev Microbiol 12(9):635–645. doi:10.1038/nrmicro3330.

3. Vandamme P, Gillis M, De Vos P, Kersters K, Swings J (1996) Polyphasic Taxonomy, a Consensus Approach to Bacterial Systematics. Microbiol Rev 60(2):407–438.

4. Patin N V, Duncan KR, Dorrestein PC, Jensen PR (2016) Competitive Strategies Differentiate Closely Related Species of Marine Actinobacteria. ISME J 10(2):478–490. doi:10.1038/ismej.2015.128.

5. Ziemert N, et al. (2014) Diversity and Evolution of Secondary Metabolism in the Marine Actinomycete Genus Salinispora. Proc Natl Acad Sci 111(12):E1130–E1139. doi:10.1073/pnas.1324161111.

6. Penn K, et al. (2009) Genomic Islands Link Secondary Metabolism to Functional Adaptation in Marine Actinobacteria. ISME J 3(10):1193–1203. doi:10.1038/ismej.2009.58.

7. Jensen PR (2016) Natural Products and the Gene Cluster Revolution. Trends Microbiol 24(12):968–977. doi:10.1016/j.tim.2016.07.006.

8. Karas M, Hillenkamp F (1988) Laser Desorption Ionization of Proteins with Molecular Masses Exceeding 10,000 Daltons. Anal Chem 60(20):2299–2301. doi:10.1021/ac00171a028.

9. Tanaka K, et al. (1988) Protein and Polymer Analyses up to m/z 100 000 by Laser Ionization Time-of-Flight Mass Spectrometry. Rapid Commun Mass Spectrom 2(8): 151–153. doi:10.1002/rcm.1290020802.

10. Sandrin TR, Goldstein JE, Schumaker S (2013) MALDI TOF MS Profiling of Bacteria at the Strain Level: A Review. Mass Spectrom Rev 32(3):188–217. doi:10.1002/mas.21359.

11. Cain TC, Lubman DM, Weber WJ, Vertes A (1994) Differentiation of Bacteria Using Protein Profiles from Matrix-Assisted Laser Desorption/Ionization Time-of-Flight Mass Spectrometry. Rapid Commun Mass Spectrom 8(12): 1026–1030. doi:10.1002/rcm.1290081224.

12. Holland RD, et al. (1996) Rapid Identification of Intact Whole Bacteria Based on Spectral Patterns Using Matrix-Assisted Laser Desorption/Ionization with Time-of-flight Mass Spectrometry. Rapid Commun Mass Spectrom 10(10):1227–1232. doi:10.1002/(SICI)1097-0231(19960731)10:10<1227::AID-RCM659>3.0.CO;2-6.

13. Maier T, Klepel S, Renner U, Kostrzewa M (2006) Fast and Reliable MALDI-TOF MS-Based Microorganism Identification. Nat Methods 3(4):68–71. doi:10.1038/nmeth870.

14. Dubois D, et al. (2012) Performances of the Vitek MS Matrix-Assisted Laser Desorption Ionization-Time of Flight Mass Spectrometry System for Rapid Identification of Bacteria in Routine Clinical Microbiology. J Clin Microbiol 50(8):2568–2576. doi:10.1128/JCM.00343-12.

15. Rahi P, Prakash O, Shouche YS (2016) Matrix-Assisted Laser Desorption/Ionization Time-of-Flight Mass-Spectrometry (MALDI-TOF MS) Based Microbial Identifications: Challenges and Scopes for Microbial Ecologists. Front Microbiol 7. doi:10.3389/fmicb.2016.01359.

16. Popović NT, Kazazić SP, Strunjak-Perović I, Čož-Rakovac R (2017) Differentiation of Environmental Aquatic Bacterial Isolates by MALDI-TOF MS. Environ Res 152:7–16. doi:10.1016/j.envres.2016.09.020.

17. Munteanu B, Hopf C (2013) Emergence of Whole-Cell MALDI-MS Biotyping for High-Throughput Bioanalysis of Mammalian Cells? Bioanalysis 5(8):885–893. doi:10.4155/bio.13.47.

18. Teramoto K, et al. (2007) Phylogenetic Classification of Pseudomonas putida Strains by MALDI-MS Using Ribosomal Subunit Proteins as Biomarkers. Anal Chem 79(22):8712–8719. doi:10.1021/AC701905R.

19. Arnold RJ, Reilly JP (1999) Observation of Escherichia coli Ribosomal Proteins and Their Post translational Modifications by Mass Spectrometry. Anal Biochem 269(1): 105–112. doi:10.1006/abio.1998.3077.

20. Silva R, Lopes NP, Silva DB (2016) Application of MALDI Mass Spectrometry in Natural Products Analysis. Planta Med 82:671–689. doi:10.1055/s-0042-104800.

21. Zeigler DR, et al. (2008) The origins of 168, W23, and other Bacillus subtilis legacy strains. J Bacteriol 190(21):6983–6995 doi:10.1128/JB.00722-08.

22. Schroeder JW, Simmons LA (2013) Complete Genome Sequence of Bacillus subtilis Strain PY79. Genome Announc 1(6). doi:10.1128/genomeA.01085-13.

23. Stein T (2005) Bacillus subtilis Antibiotics: Structures, Syntheses and Specific Functions. Mol Microbiol 56(4):845–857. doi:10.1111/j.1365-2958.2005.04587.x.

24. Candela T, Fouet A (2006) Poly-Gamma-Glutamate in Bacteria. Mol Microbiol 60(5):1091–1098.

25. Yang Y-L, et al. (2011) Connecting Chemotypes and Phenotypes of Cultured Marine Microbial Assemblages by Imaging Mass Spectrometry. Angew Chemie Int Ed 50(26):5839–5842. doi:10.1002/anie.201101225.

26. Yoon S-H, et al. (2017) Introducing EzBioCloud: A Taxonomically United Database of 16S rRNA Gene Sequences and Whole-Genome Assemblies. Int J Syst Evol Microbiol 67(5):1613–1617. doi:10.1099/ijsem.0.001755.

27. Wang M, et al. (2016) Sharing and Community Curation of Mass Spectrometry Data with Global Natural Products Social Molecular Networking. Nat Biotechnol 34(8):828–837. doi:10.1038/nbt.3597.

28. Gowda H, et al. (2014) Interactive XCMS Online: Simplifying Advanced Metabolomic Data Processing and Subsequent Statistical Analyses. Anal Chem 86(14):6931–6939. doi:10.1021/ac500734c.

29. Arias AA, et al. (2015) Growth of Desferrioxamine-Deficient Streptomyces Mutants through Xenosiderophore Piracy of Airborne Fungal Contaminations. FEMS Microbiol Ecol (7). doi:10.1093/femsec/fiv080.

30. D’Onofrio A, et al. (2010) Siderophores from Neighboring Organisms Promote the Growth of Uncultured Bacteria. Chem Biol 17(3):254–264. doi:10.1016/j.chembiol.2010.02.010.

31. Bruns H, et al. (2017) Function-Related Replacement of Bacterial Siderophore Pathways. ISME J. doi:10.1038/ismej.2017.137.

32. Blondel VD, Guillaume J-L, Lambiotte R, Lefebvre E (2008) Fast Unfolding of Communities in Large Networks. J Stat Mech Theory Exp (10). doi:10.1088/1742-5468/2008/10/P10008.

33. Gonzalez DJ, et al. (2012) Observing the Invisible through Imaging Mass Spectrometry, a Window Into the Metabolic Exchange Patterns of Microbes. J Proteomics 75(16):5069–5076. doi:10.1016/j.jprot.2012.05.036.

34. Yang Y-L, Xu Y, Straight P, Dorrestein PC (2009) Translating Metabolic Exchange With Imaging Mass Spectrometry. Nat Chem Biol 5(12):885–887. doi:10.1038/nchembio.252.

35. Bastian M, Heymann S, Jacomy M (2009) Gephi: An Open Source Software for Exploring and Manipulating Networks Visualization and Exploration of Large Graphs. Int AAAI Conf Weblogs Soc Media:361–362. doi:10.13140/2.1.1341.1520.

36. Yang JY, et al. (2013) Molecular Networking as a Dereplication Strategy. J Nat Prod 76(9):1686–1699. doi:10.1021/np400413s.

37. Zarins-Tutt JS, et al. (2016) Prospecting for New Bacterial Metabolites: A Glossary of Approaches for Inducing, Activating and Upregulating the Biosynthesis of Bacterial Cryptic or Silent Natural Products. Nat Prod Rep 33(1):54–72. doi:10.1039/c5np00111k.

38. Freiwald A, Sauer S (2009) Phylogenetic Classification and Identification of Bacteria by Mass Spectrometry. Nat Protoc 4(5):732–742. doi:10.1038/nprot.2009.37.

39. Schumann P, Maier T (2014) MALDI-TOF Mass Spectrometry Applied to Classification and Identification of Bacteria. Methods Microbiol 41:275–306. doi:10.1016/bs.mim.2014.06.002.

40. Pruesse E, Peplies J, Glöckner FO (2012) SINA: Accurate High-Throughput Multiple Sequence Alignment of Ribosomal RNA Genes. Bioinformatics 28(14):1823–1829. doi:10.1093/bioinformatics/bts252.

41. Bligh EG, Dyer WJ (1959) A Rapid Method of Total Lipid Extraction and Purification. Can J Biochem Physiol 37(8):911–917. doi:10.1139/o59-099.

42. Goering AW, et al. (2016) Metabologenomics: Correlation of Microbial Gene Clusters with Metabolites Drives Discovery of a Nonribosomal Peptide with an Unusual Amino Acid Monomer. ACS Cent Sci 2(2):99–108. doi:10.1021/acscentsci.5b00331.

43. Tautenhahn R, Patti GJ, Rinehart D, Siuzdak G (2012) XCMS Online: A Web-Based Platform to Process Untargeted Metabolomic Data. Anal Chem 84(11):5035–5039. doi:10.1021/ac300698c.

44. Martens L, et al. (2011) mzML--A Community Standard for Mass Spectrometry Data. Mol Cell Proteomics (1). doi:10.1074/mcp.R110.000133.

45. Chambers MC, et al. (2012) A Cross-Platform Toolkit for Mass Spectrometry and Proteomics. Nat Biotechnol 30(10):918–920. doi:10.1038/nbt.2377.

46. Gibb S, Strimmer K (2012) MALDIquant: A Versatile R Package for the Analysis of Mass Spectrometry Data. Bioinformatics 28(17):2270–2271. doi:10.1093/bioinformatics/bts447.

47. Chang W, Cheng J, Allaire J, Xie Y, McPherson J (2016) Shiny: Web Application Framework for R.

